# Imputing Single-Cell Protein Abundance in Multiplex Tissue Imaging

**DOI:** 10.1101/2023.12.05.570058

**Authors:** Raphael Kirchgaessner, Cameron Watson, Allison Creason, Kaya Keutler, Jeremy Goecks

## Abstract

Multiplex tissue imaging are a collection of increasingly popular single-cell spatial proteomics and transcriptomics assays for characterizing biological tissues both compositionally and spatially. However, several technical issues limit the utility of multiplex tissue imaging, including the limited number of molecules (proteins and RNAs) that can be assayed, tissue loss, and protein probe failure. In this work, we demonstrate how machine learning methods can address these limitations by imputing protein abundance at the single-cell level using multiplex tissue imaging datasets from a breast cancer cohort. We first compared machine learning methods’ strengths and weaknesses for imputing single-cell protein abundance. Machine learning methods used in this work include regularized linear regression, gradient-boosted regression trees, and deep learning autoencoders. We also incorporated cellular spatial information to improve imputation performance. Using machine learning, single-cell protein expression can be imputed with mean absolute error ranging between 0.05-0.3 on a [0,1] scale. Finally, we used imputed data to predict whether single cells were more likely to come from pre-treatment or post-treatment biopsies. Our results demonstrate (1) the feasibility of imputing single-cell abundance levels for many proteins using machine learning; (2) how including cellular spatial information can substantially enhance imputation results; and (3) the use of single-cell protein abundance levels in a use case to demonstrate biological relevance.

## Introduction

Multiplex tissue imaging (MTI) are a set of single-cell spatial proteomics and transcriptomics assays for highly detailed profiling of biological tissues. With MTI, single-cell abundance levels and spatial distribution of 10-150 of proteins and/or 500-2000 RNAs can be quantified simultaneously (Francisco-Cruz et al., 2020; Sheng et al., 2023). MTI enables characterization of individual cells as well as tissue organization, and MTI has been used in studies of healthy tissue (Neumann et al., 2022), COVID (Werlein et al., 2023), cancer (Lewis et al., 2021), and other diseases (Kitko et al., 2022; McCaffrey et al., 2022; Sepe et al., 2022). There are many MTI platforms, including cyclic immunofluorescence (CycIF) (Lin et al., 2018), CO-Detection by indEXing (CODEX) (Black et al., 2021), CosMx (Z. R. Lewis et al., 2022), Xenium (Janesick et al., 2022) and multiplex immunohistochemistry (Tsujikawa et al., 2017). MTI has been used to generate large datasets in NIH consortia such as the NIH Human BioMolecular Atlas Program (HuBMAP Consortium, 2019) and the NCI Cancer Moonshot Human Tumor Atlas Network (Rozenblatt-Rosen et al., 2020). MTI is also an increasingly common assay in cancer (Tan et al., 2020), where it has proven important for quantifying tumor spatial organization and microenvironment heterogeneity (Blise et al., 2022) and connecting these features to cancer subtypes, prognosis, and therapy response (Friebel et al., 2020; Steele et al., 2020).

However, several key factors limit the usefulness of MTI. Only 10-150 proteins and/or several thousand RNAs can be assayed in a single experiment, and hence the information obtained from a single experiment is bounded. Further, MTI assays can suffer from several technical issues that reduce the information obtained, including tissue loss or folding, probe failure, illumination artifacts, or errors in downstream image processing. These limitations greatly impact MTI data quality and substantially reduce the overall utility of MTI. To mitigate these limitations and improve utility of MTI, machine learning and deep learning approaches can be used to computationally increase the numbers of proteins/RNAs available from MTI and mitigate assay failures. Computationally increasing—or imputing—additional data by filling in missing data with predicted values is already common in other molecular assays, such as single-cell RNA sequencing (scRNA) (Chen et al., 2022; Gong et al., 2018; He et al., 2020; Kharchenko et al., 2014; Talwar et al., 2018; Tran et al., 2022; van Dijk et al., 2018; Xu et al., 2021), bulk genomics (Qiu et al., 2020), and bulk transcriptomics (Patruno et al., 2021). While imputation has been applied to MTI images (Sims & Chang, 2023; Ternes et al., 2021), to the best of our knowledge imputation on MTI single-cell datasets has not been explored. Imputation has been applied to MTI image data (Pati et al., 2023; Wu et al., 2023), being able to reconstruct protein expression in images. However, imputing single-cell data is especially valuable because single-cell datasets require fewer computational resources to process than images and can be readily integrated with other molecular datasets.

In this study, we applied machine learning (ML) and deep learning (DL) methods to impute protein abundance in tissue-based cyclic immunofluorescence (t-CyCIF) (Lin et al., 2018) datasets obtained from breast cancer tissues. Because t-CyCIF is an open and quantitative multiplexed tissue imaging assay, it is ideally suited for imputation. We evaluated the performance of ML and DL methods to predict protein abundance levels in t-CyCIF single-cell datasets that included 20 proteins. Three distinct ML/DL approaches—regularized linear regression, gradient-boosted trees, and autoencoders—were used to impute single-cell protein abundance values across both patients and timepoints. Spatial information was introduced to improve imputation results. To demonstrate a biological application of imputed single-cell protein abundance, we used imputed data to predict whether single cells were more likely to come from pre-treatment or post-treatment breast cancer biopsies Overall, our results demonstrate that accurate imputation is possible for many proteins, that spatial information significantly improves imputation results, and that imputed protein values are useful in a biological application.

## Results

### Study Cohort and Analysis Overview

The multiplexed tissue imaging single-cell datasets used in this study were generated using a 20-plex t-CyCIF (Lin et al., 2018) protein panel applied to a cohort of hormone receptor-positive (HR+), HER-2 negative metastatic breast cancer biopsies. t-CyCIF is a unique multiplexed tissue imaging assay that has been shown to provide robust and repeatable quantifications of protein concentrations across a range of biological samples. The tissue biopsies and datasets are part of the NCI Cancer Moonshot Human Tumor Atlas Network (Rozenblatt-Rosen et al., 2020) and have detailed associated clinical metadata. Our dataset includes a total of eight biopsies derived from four patients (**Fig. 1a**) that received a CDK4/6 inhibitor in combination with endocrine therapy, which is a common combination therapy in metastatic HR+ breast cancer. Each patient contributed a pair of biopsies, a pre-treatment biopsy and a biopsy taken at the time of tumor progression.

**Figure 1:**
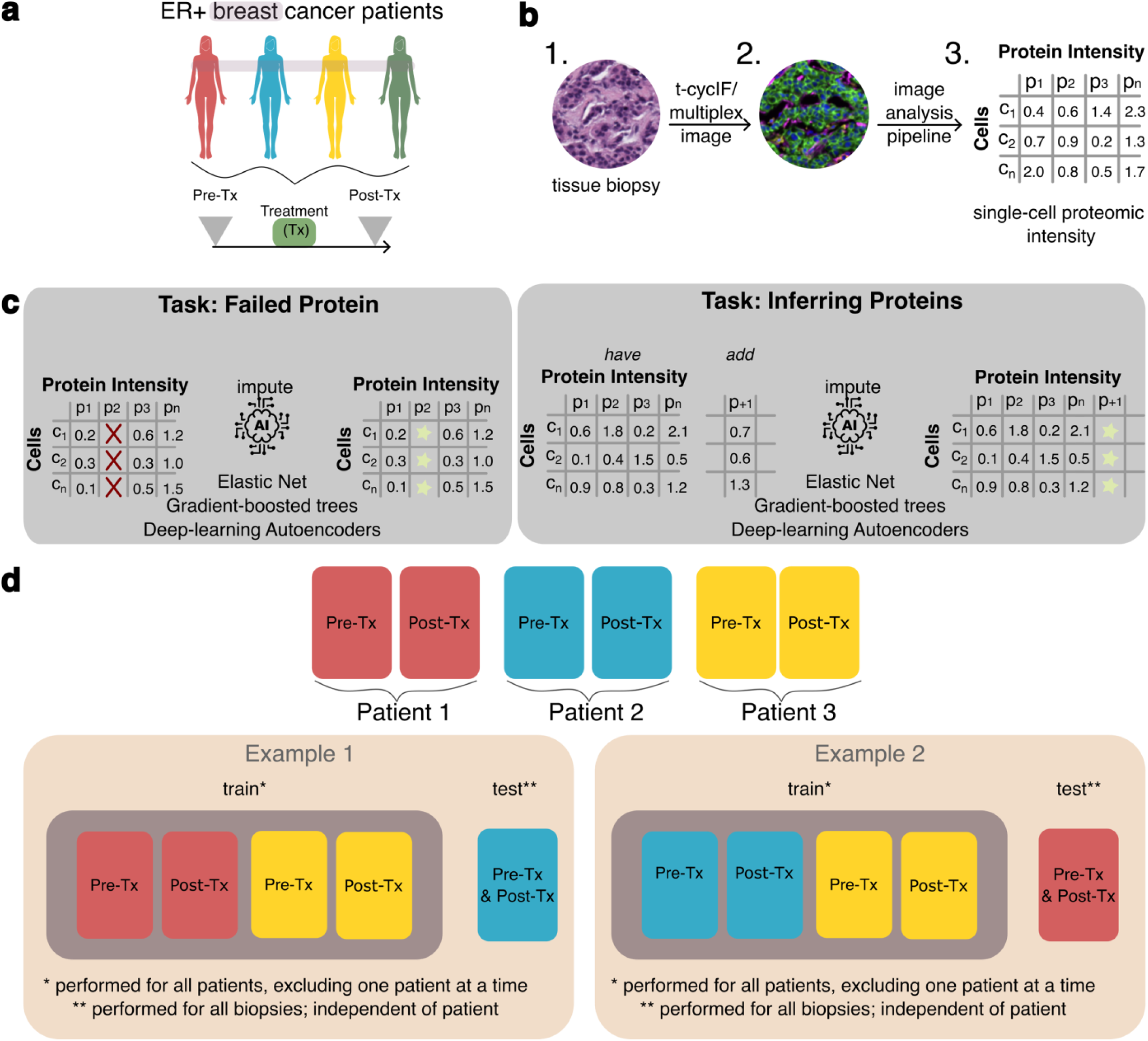
Overview of dataset, study motivations, and analysis approaches. **a:** Biopsies were obtained from four HR+ breast cancer patients before and after the same standard-of-care therapy for a total of eight biopsies. **b:** each biopsy was assayed using the multiplexed tissue imaging assay t-CyCIF to quantify abundance levels of 20 proteins and then processed using an image analysis pipeline to create single-cell feature tables (total number of cells identified: 475359); **c:** the key tasks addressed by this work are imputing failed proteins and inferring additional proteins not present in an multiplex tissue imaging (MTI) experiment; **d:** approaches for training and testing ML models for imputing proteins across patients.

Image stacks collected from t-CyCIF were processed using the MCMICRO image analysis pipeline (Schapiro et al., 2021) to generate single-cell feature tables (**Fig. 1b**). Each row in the table is a single cell identified in the image, and the table columns are the protein abundance levels calculated via mean pixel intensity per cell. In total 475,359 single cells were identified across all biopsies, with an average of 59,400 cells per biopsy. To perform in-patient evaluation, either the pre-treatment or the post-treatment biopsy was used for training a machine learning model while the remaining biopsies were used for testing model performance. Biopsy timing was not used in this study. In total 16 proteins were shared between all biopsies, including eight proteins for identifying cell types (lineage proteins) and eight proteins for characterizing cellular functional states (functional proteins) (**Table 1**).

**Table 1:** Overview of proteins assayed using t-CyCIF and their use as functional or lineage proteins. Lineage proteins are used to identify cell types whereas functional proteins are used to characterize cell function.

| Lineage Protein | Functional Protein |
| --- | --- |
| CD45 | Ki67 |
| αSMA | pERK |
| eCadherin | PR |
| CK19 | EGFR |
| CK14 | p21 |
| CK17 | pRB |
| CK7 | AR |
| Vimentin | HER2 |

The imputation task in this study was to predict protein abundance levels for a withheld protein or set of proteins. Preprocessing was performed to remove all columns from the datasets except protein intensities, followed by a min-max scaling approach, which maps values between zero and one. Thus, model error is in the range [0,1] where lower error represents better performance. For each machine learning experiment, one or more proteins were withheld and used as the target variable(s) for the predictive model, and the remaining protein abundances were used as input features for the model. This task simulates the key application for imputation in MTI: computationally increasing proteins not originally included in an MTI assay or inferring protein levels where the assay failed (**Fig. 1c**). Three machine learning methods were used for imputation: elastic-net (EN) regularized linear regression (Su et al., 2012), light gradient-boosting machine (LGBM) (Ke et al., 2017), and neural network autoencoders (AE) (Zhai et al., 2018).

These algorithms offer different advantages for addressing the complexities of our dataset and research objectives. Elastic Net (EN) is a linear model that effectively handles high-dimensional data like that in our MTI datasets by using regularization to manage multicollinearity and select relevant proteins. However, EN requires a separate model to predict each protein, and this is time-consuming. Light Gradient Boosting Machines (LGBM) use a non-linear approach for developing predictive models and are amongst the most efficient and best performing methods for tabular data like our MTI datasets. Like EN, LGBM also requires a model for each protein. Autoencoders (AEs) can learn non-linear relationships, reduce dimensionality, and denoise data, allowing a single model to impute multiple proteins at once, although their compression techniques may lead to some loss of precision. Autoencoders were chosen for this study due to their ability to reduce dimensionality and denoise data while preserving essential information. This helps in data imputation and improving data quality before further analysis. EN and LGBM are straightforward to implement and handle linear and non-linear relationships, respectively, while AEs provide efficient preprocessing and multi-protein imputation.

Imputation model training and evaluation were conducted using a leave-one-out cross-validation (LOOCV) approach (**Fig. 1d**). In this methodology, each patient was considered a single data point, whereby a model was trained on all biopsies except those from one patient. Subsequently, the model’s performance was assessed using the biopsies from the patient excluded during training. This LOOCV approach was chosen to prevent data leakage from biopsies associated with the same patient as the test biopsy, thereby closely approximating real-world application scenarios. Model performance was calculated by averaging the mean absolute error (MAE) scores across all runs of the model on a particular train-test dataset split. Statistical evaluations were carried out using the Mann-Whitney U test, and multiple hypothesis tests were adjusted using the Benjamini-Hochberg correction method.

### Protein abundance imputation with elastic net and light gradient-boosting machines

To establish baseline performance of our imputation models, we conducted a test using mean imputation, where values were imputed by using the mean protein abundance value in the training dataset. Using mean values for imputation serves as a null or baseline model to determine if a machine learning model provides genuine improvements over a simple heuristic. The EN model outperformed the null model by an average of 0.078 MAE indicating that the EN model demonstrated superior performance compared to the null model (**Fig. 2a**). This performance difference was statistically significant, with an average adjusted p-value less than 0.0001 for all proteins. Proteins CK17 and Ki67 were most accurately imputed with MAE of 0.05. Proteins for which the imputation MAE exceeded 0.2 included CK19, ER, CK14, and PR.

**Figure 2:**
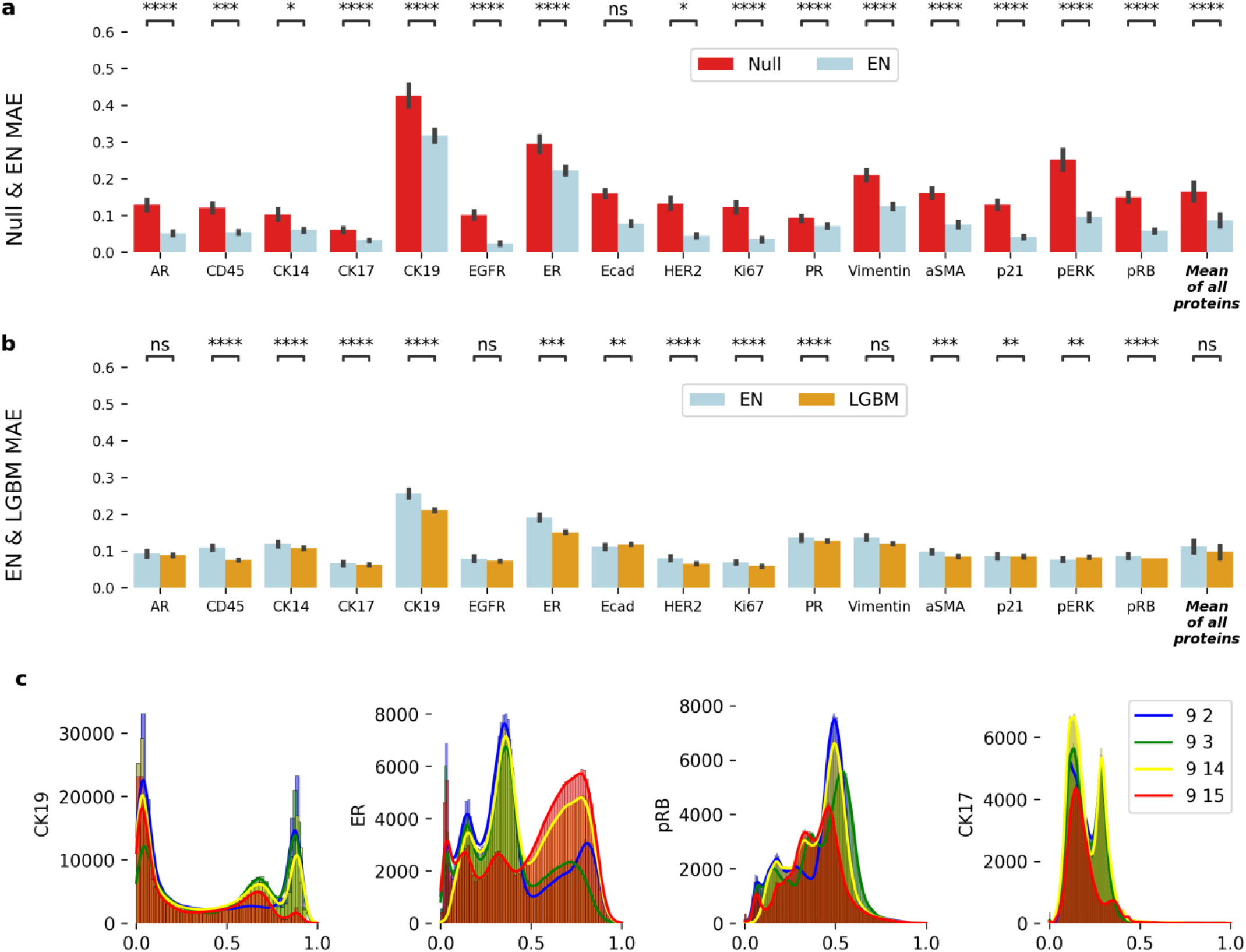
Imputation results for null model and elastic net and Light GBM machine learning models across patients. **a:** Imputation results for all proteins demonstrates improved mean absolute error (MAE) by using Elastic Net (EN) compared to a null model. **b:** Imputation results using EN & Light GBM (LGBM) show low MAE for imputation for 12 out of 16 available proteins. **c**: Patient protein expression for four proteins (CK19, ER, pRB, CK17) is highly variable; p-value: ns: not significant, p <= 1.00e+00 *: 1.00e-02 < p <= 5.00e-02 **: 1.00e-03 < p <= 1.00e-02 ***: 1.00e-04 < p <= 1.00e-03 ****: p <= 1.00e-0

Using Light Gradient Boosting Machine (LGBM) yielded improved imputation accuracy compared to EN (**Fig. 2b**). Like the EN, LGBM performance for the same 12 of 16 proteins was between 0.05 and 0.20 MAE. LGBM imputation accuracy for CK19 and ER are like the EN and greater than 0.2 MAE. Overall, LGBM displayed more accurate imputation results than EN (**Table 2**) both in terms of mean and standard deviation. To provide a comprehensive overview of performance, a ***mean of all proteins*** column is included to show the average imputation accuracy across all proteins for each model (**Fig. 2a, Fig. 2b**). While using LGBM improved imputation accuracy compared to the EN, some proteins still exhibit a high MAE, such as CK19 and ER. These proteins exhibited high variance (**Table S2**), presenting a significant challenge for accurate imputation. **Figure 2c** shows protein abundance distributions of selected proteins with especially high or low variance to illustrate why imputation is difficult for proteins such as CK19 and ER that exhibit high variance.

**Table 2:** Mean and standard deviation of performance for EN, LGBM and AE.

| Model/Network | EN | LGBM | AE Single | AE Multi |
| --- | --- | --- | --- | --- |
| Mean (Std) | 0.11 (0.07) | 0.10 (0.06) | 0.13 (0.09) | 0.13 (0.09) |

We also evaluated imputation accuracy within patients by modifying the LOOCV approach described above. The modified within-patients LOOCV approach included one biopsy from each patient in the training dataset and used the remaining biopsy from the same patient for testing. Surprisingly, imputation accuracy in the across-patient LOOCV approach was higher than imputation accuracy in the within patients for some proteins (**Fig. S1**). We hypothesize that this performance difference may be attributable to the more diverse training dataset in the across-patient approach. This diverse training dataset may enable imputation models to better handle heterogeneity across patients.

### Protein abundance imputation using autoencoders and all model comparisons

An autoencoder (AE) is a deep learning neural network for accurate reconstruction of high-dimensional data that include two distinct components: (1) an encoder network that maps a high-dimensional input to a lower-dimensional representation in a latent space and (2) a decoder network that reconstructs the original high-dimensional input from the low-dimensional latent space representation. The goal of an AE is to perform information-preserving dimensionality reduction of its input to the latent space so that it can then accurately reconstruct the input from the latent representation. AEs have been successfully used for imputation in various biological domains, including single-cell RNA (Grønbech et al., 2020; Hou et al., 2020; Lotfollahi et al., 2020; Talwar et al., 2018), genomics (Qiu et al., 2020) and more (McCoy et al., 2018). Unlike LGBM and EN models, AEs can impute multiple features simultaneously due to their ability to fully reconstruct the entire input data. Leveraging this capability, we conducted both single-protein and multi-protein imputation experiments based on the order of protein assays during t-CyCIF’s multiple imaging rounds. T-CyCIF involves multiple rounds to stain, incubate, and capture images. Proteins were sequentially removed from each round, and the AE was trained and evaluated for each set of proteins. Initially, proteins from the first round were removed, and the AE was trained and evaluated. This process was then repeated for all proteins in the second round, and so on. To maintain simplicity, no other pairings of proteins were made beyond the rounds.

The AE was trained using biopsies from three patients, including both pre- and post-treatment samples, with biopsies from a fourth patient reserved for validation. Aggregating all biopsy data allowed the AE to develop an internal representation focused on minimizing reconstruction error. During the imputation phase, we initially replaced the target protein’s values with the mean expression levels across the dataset. The modified dataset was processed through the AE, which performed continuous cycles of encoding and decoding to iteratively refine the imputed values. For each cycle, the AE replaced the original protein values with the decoded data from the previous cycle. This iterative process was repeated 10 times, as each protein required a different number of optimal iterations for accurate imputation. We used the mean expression from iterations five to ten as the final imputed value (**Fig. 3a**).

**Figure 3:**
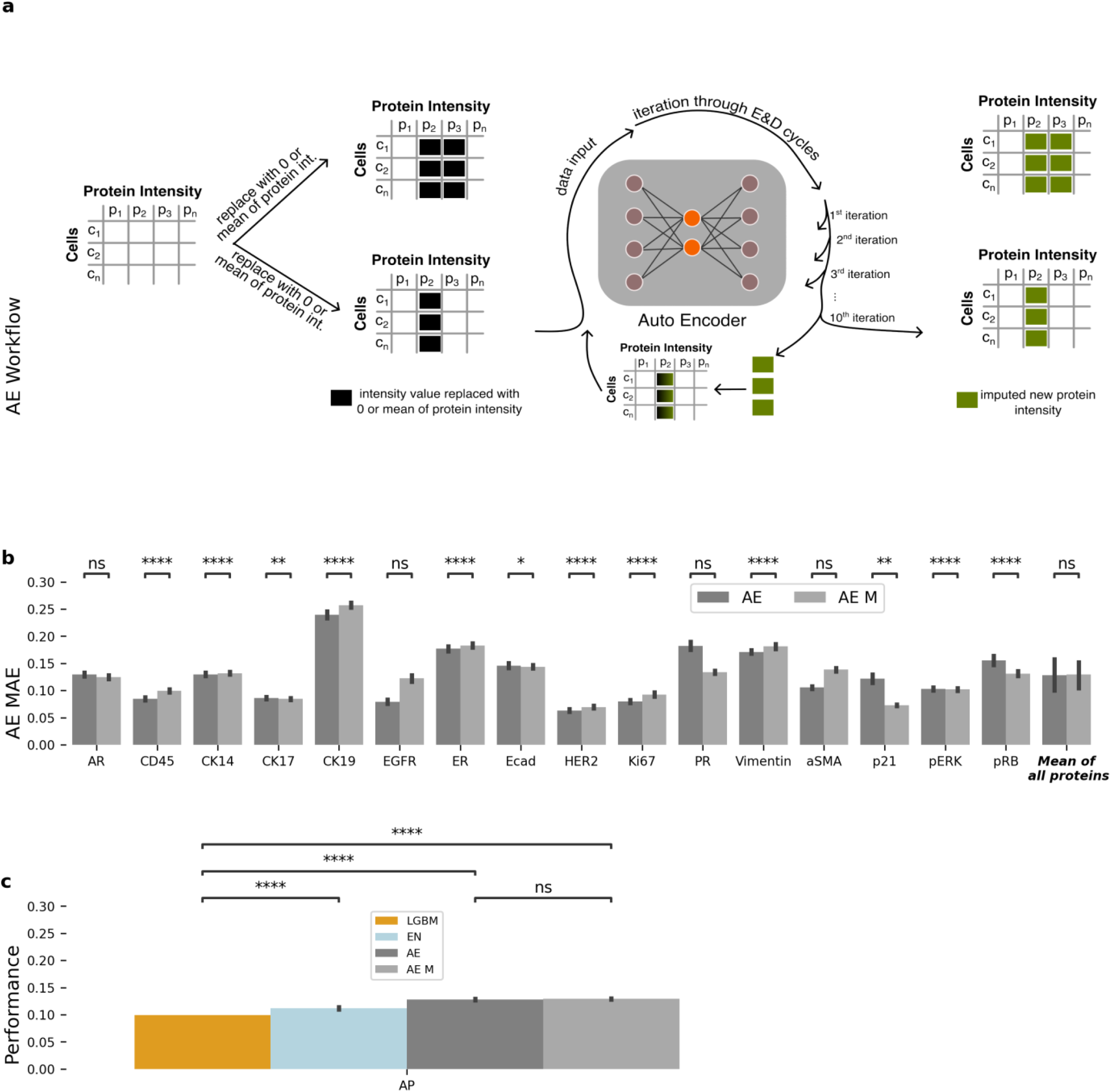
Autoencoder imputation results and performance comparison between machine learning models. **a**: The autoencoder (AE) is trained and then uses an iterative approach to impute single or multiple proteins. To start, proteins to be imputed are replaced with either zero or the mean of the intensity values in the training set. Then, the autoencoder is used iteratively to predict protein intensities using output values as new input values for each iteration. **b**: AE single- and multi-protein imputation performance. **c**: performance comparison between all evaluated ML models shows similar performance overall and that LGBM performs best, followed by EN and finally AE. There is no significant difference between single and multi-protein imputation performance for AE. p-values: ns: not significant p <= 1.00e+00 *: 1.00e-02 < p <= 5.00e-02 **: 1.00e-03 < p <= 1.00e-02 ***: 1.00e-04 < p <= 1.00e-03 ****: p <= 1.00e-0

AEs accurately imputed proteins in both single- and multi-protein experiments (**Fig. 3b**). Imputation accuracy of CK19 levels is between 0.15-0.35 MAE, while imputation of the best performing proteins, CK17 and p21, is between 0.05 and 0.10 MAE. Like the EN and LGBM models, imputation performance is worst for the proteins with the most variable abundance levels in our breast cancer cohort, including CK19, ER, and PR.

We next compared performance for all three machine learning models used for imputation. Overall, LGBM performed best, followed by the EN and the AEs. These performance differences are consistent between models (**Fig. 3c**). However, performance differences between the models are relatively modest, with the LGBM achieving a mean accuracy of 0.10 MAE, followed by the EN with a mean accuracy of 0.11 MAE, and the AEs with a mean accuracy of 0.13 MAE (**Fig. 3c**). Autoencoder performance differences between the imputation of single proteins and that of multi-proteins are minimal.

To further evaluate the performance of imputation in MTI using machine learning, we performed imputation on an additional t-CyCIF dataset from the Human Tumor Atlas Network (Rozenblatt-Rosen et al., 2020). This dataset was taken from a breast cancer tissue microarray and included two tissue cores from each of 26 breast cancer tumors. On average, each core included approximately 9850 cells.

Unlike the cancers in our main analysis cohort, these cancers are primary disease rather than metastatic and represent all different subtypes of breast cancer. This data is publicly available at the NCI Human Tumor Atlas Portal (see Data Availability section), and the same primary image processing analysis pipeline used in our main analysis was used to generate a single-cell dataset for imputation. From this single-cell dataset of a breast cancer tissue microarray, we extracted the proteins shared with the original dataset. We then replicated our original imputation experiments using the EN, LGBM, and AE models, employing the same LOOCV approach as before. Results from this analysis show accurate imputation from EN and LGBM models with approximately the same level of overall accuracy as in our main dataset. Accuracy of the AE models was significantly lower in this dataset as compared to accuracy in our main dataset, but it is unclear why this drop in performance occurred. The LGBM model outperformed all other models, while the AE performed better than the EN but worse than the LGBM (**Fig. 4, Table S2**).

**Figure 4:**
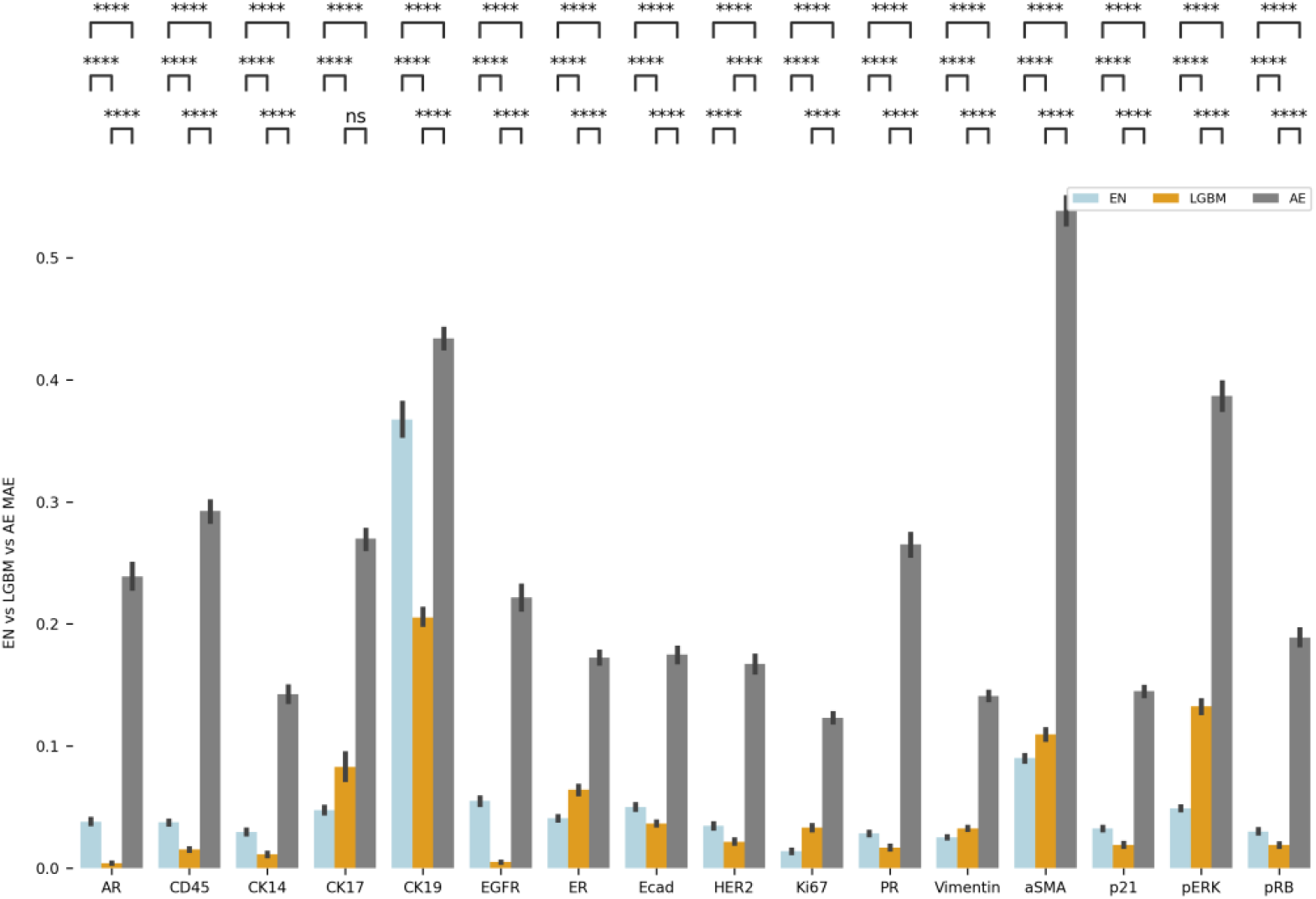
Imputation performance of EN, LGBM, and AE machine learning models on an independent t-CyCIF dataset. Dataset was obtained from a breast cancer tissue microarray that includes two cores each from 26 tumors. Imputation results are similar to those obtained in our primary cohort and dataset, showing that our imputation methods are applicable beyond the primary cohort to other cohorts and datasets. p-values: ns: p <= 1.00e+00 *: 1.00e-02 < p <= 5.00e-02 **: 1.00e-03 < p <= 1.00e-02 ***: 1.00e-04 < p <= 1.00e-03 ****: p <= 1.00e-0

### Using cellular spatial information to improve imputation

A key advantage of MTI datasets is that the spatial coordinates of each cell are known, making it possible to quantify spatial information around individual cells. We hypothesized that the spatial information available in t-CyCIF could be used to improve imputation performance. To test this hypothesis, we quantified the spatial cellular context surrounding a target cell (cell of interest) by calculating the mean protein abundance of neighboring cells. Average abundance levels of all proteins in neighboring cells were then added to our prior set of input features to create a feature set that includes both single-cell protein abundances plus average neighbor abundances (**Fig. 5a**).

**Figure 5:**
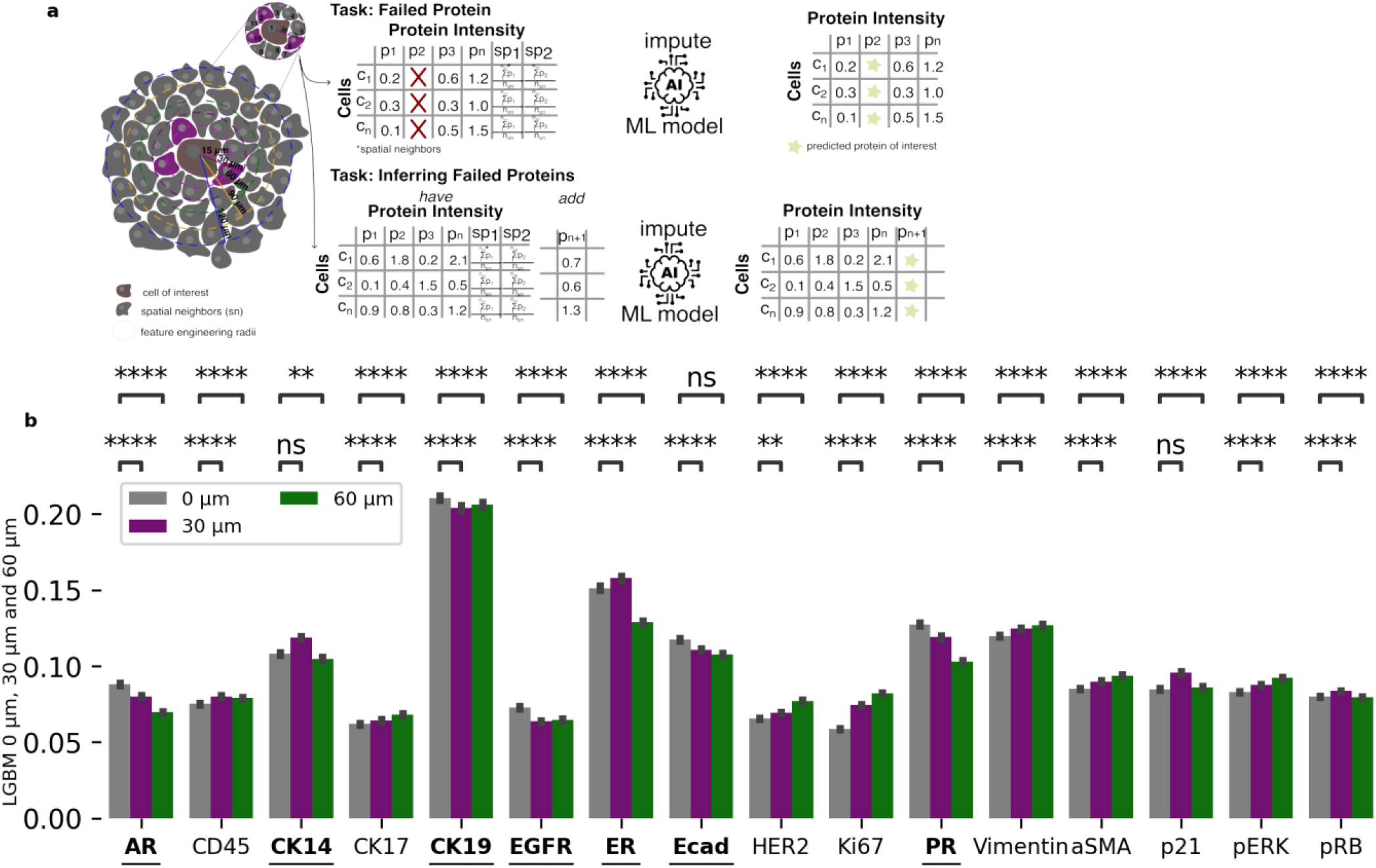
Using spatial information improves imputation performance for LGBM. **a:** Schematic for creating a feature table based on spatial neighbors found in selected radii. Exemplary 15 µm radius is shown. Red marks the cell of interest (or origin) and protein abundance levels of cells in its neighborhood are averaged to get neighborhood abundance levels. **b:** LGBM imputation results across patients with MAE scores for 0 µm, 30 µm, 60 µm reveal significant improvement for several proteins such as EGFR, ER,ECAD and PR. p-values: ns: p <= 1.00e+00 *: 1.00e-02 < p <= 5.00e-02 **: 1.00e-03 < p <= 1.00e-02 ***: 1.00e-04 < p <= 1.00e-03 ****: p <= 1.00e-04

When no neighboring cells were detected, a value of zero was assigned for neighbors’ protein abundances. Radii of 15, 30, 60, 90 and 120 micrometers (µm) were used to identify neighboring cells and assess the impact of using different sizes of radii on imputation performance. Only features for one radius setting were used for training a model, and hence a single set of spatial features was included as input for a predictive model.

A radius of 15 µm captures most of the immediate neighbors of a cell, whereas larger radii capture the extended neighborhood of a cell.

We evaluated imputation accuracy using added spatial information in only LGBM and AEs because LGBM performed better than EN and AEs can perform multi-protein imputation. Using spatial information improved overall imputation accuracy for LGBM (**Fig. 5b, Fig. S2**), single protein AE (**Fig 6a, Fig. S3**) and multi-protein AE (**Fig. 6b, Fig S4**). Importantly, imputation accuracy for proteins that had proven difficult to impute well due to their very high levels of variance (see **Supplemental Data, Fig. S3**) was improved significantly with spatial information (**Fig. 6a**). In particular, imputation of CK19, ER, and PR was much more accurate with spatial information. LGBM performance also improved for other proteins such as CK17, CD45, Ecad, ASMA and p21. Performance of the AE achieved improvements in single-protein imputation for most proteins, with CK19 showing the greatest improvement (**Fig. 6a**). Multi-protein imputation also benefited from spatial information integration (**Fig. 6b**). However, performance gains were not as pronounced compared to the single protein imputation model. Aligned with prior research (Fischer et al., 2022), imputation accuracy generally improves up to a certain neighborhood radius and then plateaus or declines (**Table 3, Fig. S1**). However, the LGBM does not show the same improvement up until a certain radius, but instead remains largely steady, with a peak performance observed using 60 µm (**Table 3, Fig. S2**). Imputation performance is improved by incorporating spatial information (**Table S1**).

**Table 3:** Mean and standard deviation of performance for LGBM and AEs for different radii. Mean and (standard deviation) for each model are listed. 0µm is considered as baseline without any use of spatial information.

| Network | 0 $\mu\text{m}$ [Baseline] | 30 $\mu\text{m}$ | 60 $\mu\text{m}$ |
| --- | --- | --- | --- |
| LGBM | 0.10 (0.06) | 0.10 (0.06) | 0.10 (0.05) |
| AE Single Protein | 0.13 (0.09) | 0.10 (0.06) | 0.11 (0.06) |
| AE Multi Protein | 0.12 (0.09) | 0.11 (0.06) | 0.12 (0.07) |

**Figure 6:**
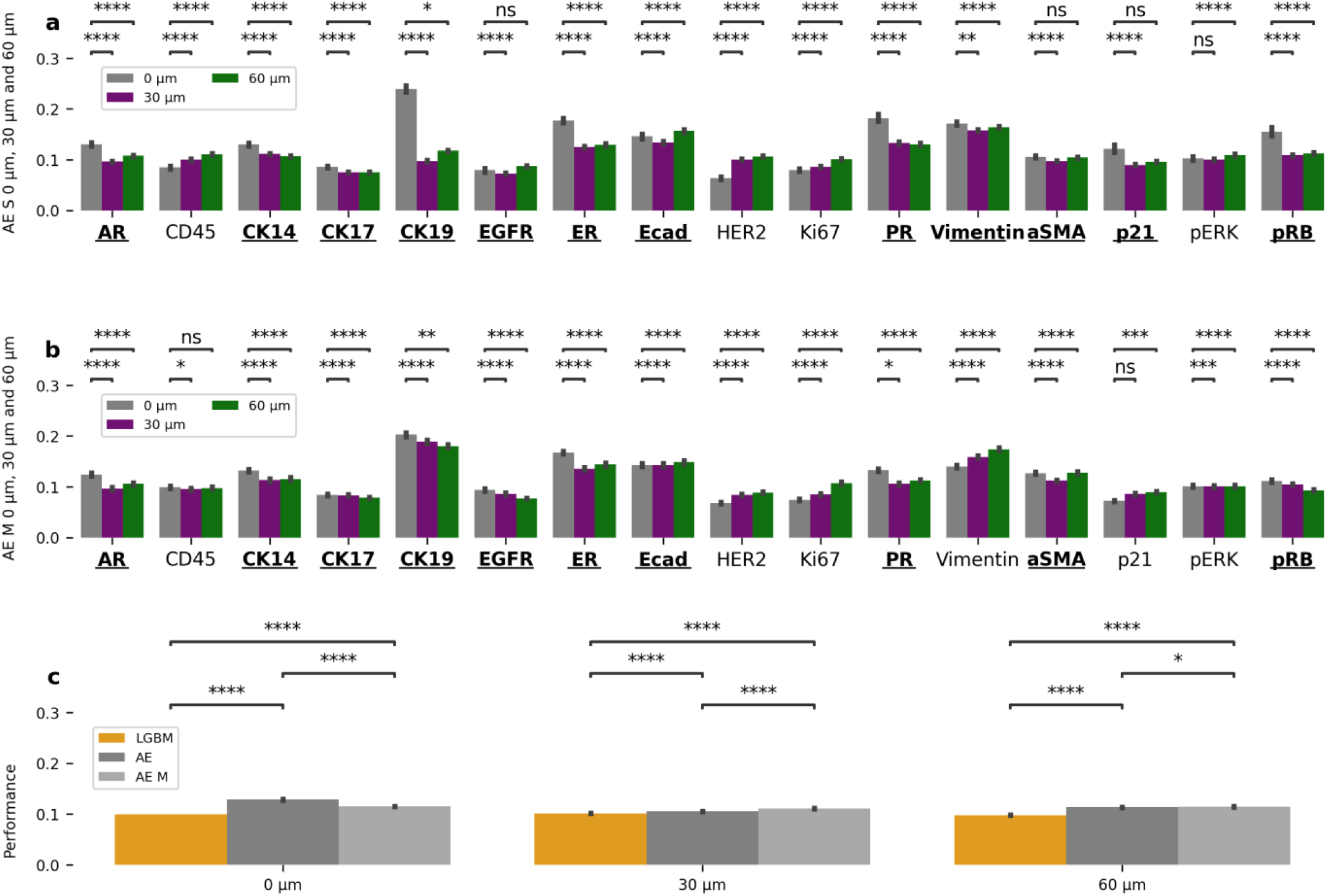
Using spatial information improves imputation performance. **a:** Single protein imputation MAE for 0 µm, 30 µm and 60 µm leads to improved imputation accuracy for proteins such as AR, CK14, CK19, ER and more. Proteins for which imputation improved when using spatial information are in bold and underlined. **b:** Multi-protein imputation MAE scores for 0 µm, 30 µm and 60 µm and leads to improved imputation accuracy for proteins such as AR, CK14, CK19, ER and more. **c:** Comparison of LGBM and AE imputation performance for 0,30 and 60µm shows similar performance of all models. p-values: ns: p <= 1.00e+00 *: 1.00e-02 < p <= 5.00e-02 **: 1.00e-03 < p <= 1.00e-02 ***: 1.00e-04 < p <= 1.00e-03 ****: p <= 1.00e-04

### Using Imputation to Predict Treatment Timepoints of Breast Cancer Cells

To evaluate the utility of imputed single-cell protein values from our machine learning models, we used these imputed values to predict whether cells were in pre-treatment or post-treatment timepoints. Using a machine learning classifier that predicts whether single cells are most likely to come from a pre-or post-treatment, we compared classifier accuracy using imputed values with classifier accuracy using original values. The dataset used for this analysis is our primary dataset that consisted of four pre-treatment and four post-treatment biopsies.

Initially poor classifier performance was observed because not all biopsy tissues exhibited strong signals associated with a treatment timepoint. To address this issue, we developed a two-step process using two machine learning classifiers **(Fig. 7a)**. First, we selected 300 µm x 300 µm tiles for each biopsy that were associated with treatment timepoint by using a tile machine learning classifier (**Fig. 7b, Fig. 7c**). The average protein expression of all cells within a tile was used as input to this tile classifier, and the tiles correctly predicted as pre-or post-treatment by the classifier were selected and used for further analysis. Second, the cells from the correctly predicted and subsequently selected tiles were then used as the dataset for single-cell imputation and then timepoint prediction using a single-cell classifier. This single-cell classifier was trained to classify single cells as either pre-treatment or post-treatment based on protein abundance levels. Accuracy of this single-cell classifier was compared using both imputed values and the original values.

**Figure 7:**
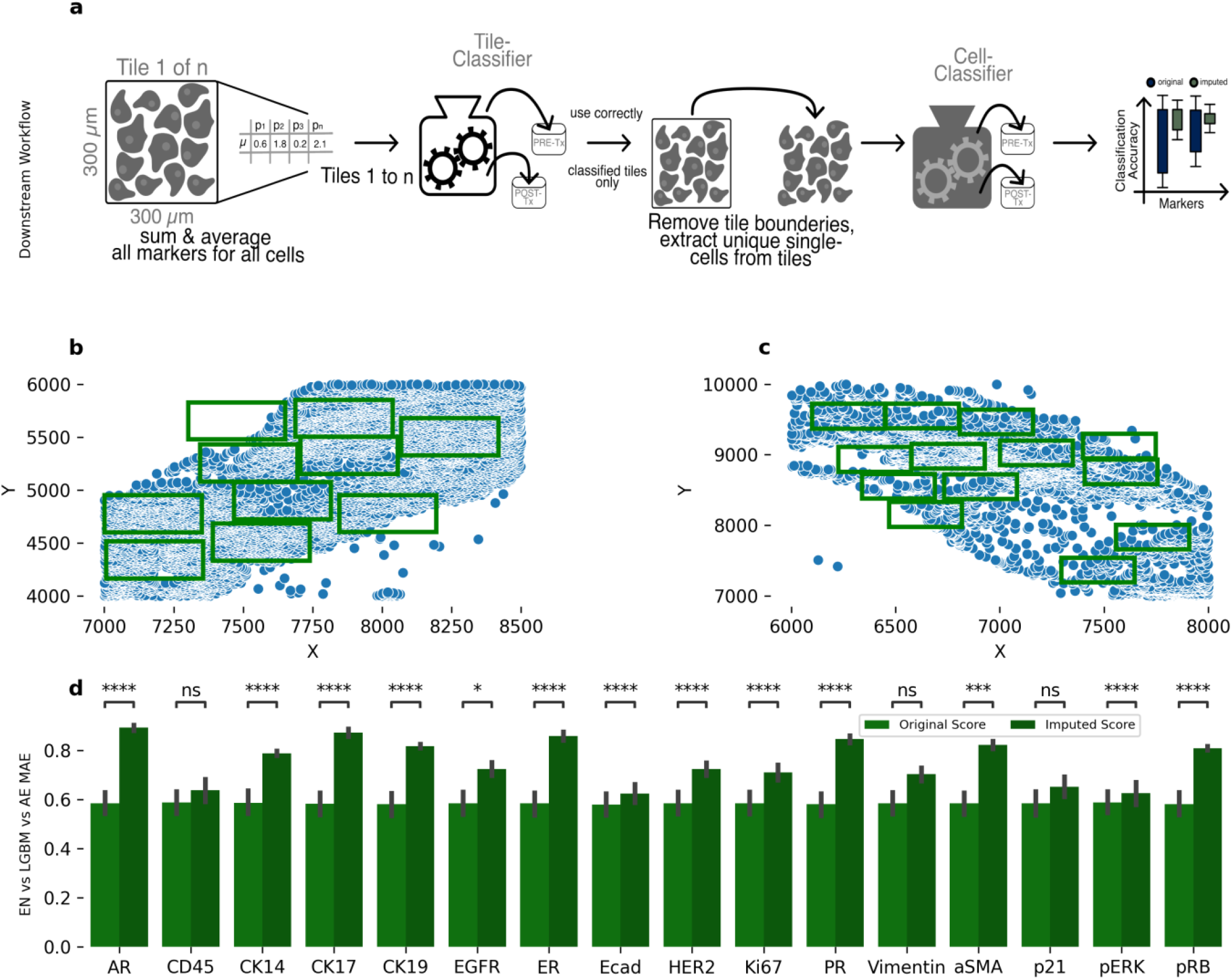
Experimental setup and validation for using imputed values to predict treatment timepoints for single cells. **a:** An initial tile classifier was used to identify tissue strongly associated with treatment timepoints. Next, cells in tissue associated with treatment timepoints were to train a cell classifier to identify whether cells came from pre-treatment or post-treatment biopsies. **b-c**: Green squares show tiles strongly associated with treatment timepoints. **d**: Classification accuracy of the cell classifier shows improved performance using imputed values as compared to performance using original values. p-values: ns: p <= 1.00e+00 *: 1.00e-02 < p <= 5.00e-02 **: 1.00e-03 < p <= 1.00e-02 ***: 1.00e-04 < p <= 1.00e-03 ****: p <= 1.00e-04

To prevent data leakage, we employed LOOCV across patients. For each iteration, a biopsy was designated as the test set, while the remaining biopsies, excluding those from the same patient, were used for training the models. We then compared classification accuracy using the original protein abundance data without adjustment to accuracy using the original data together with one protein’s values replaced with imputed data. Imputation was performed using the LGBM model because it was the most accurate in prior analyses. No spatial data was used in the imputation model for simplicity and so that a comparison between original and imputed data was straightforward. Overall, classification accuracy was higher with imputed data as compared to the original data (**Fig. 7c**). We hypothesize that the improved accuracy using the imputed data versus accuracy using the original data may result from imputation reducing the noise in the data and errors in the primary image analysis pipeline. These findings validate the biological relevance and utility of imputed data for predicting cancer treatment timepoints.

## Discussion

In this study, we utilized machine learning models to accurately impute single-cell protein abundance levels in breast cancer tissue using datasets obtained from the t-CyCIF multiplexed tissue imaging (MTI) assay. Our datasets comprised eight biopsies from a cohort of four metastatic breast cancer patients, facilitating the training and evaluation of these models. Within a range of [0,1], the imputation performance for most proteins exhibited a mean absolute error (MAE) between 0.05 and 0.15. However, proteins with high variance in our cohort, such as CK19 and ER, were more challenging to impute, with MAE ranging from 0.15 to 0.35. The LGBM model, a gradient-boosted regression tree approach, showed modestly better accuracy than an Elastic Net (EN) model or a deep learning autoencoder (AE). Incorporating spatial features into the models, represented by neighboring cells’ protein abundance levels, enhanced their accuracy and reduced the average MAE by 0.02. This improvement was particularly significant for proteins with high variance that were otherwise difficult to impute. This use of spatial information complements recent research indicating that cell communication may vary and requires careful evaluation using multiple cellular neighborhoods (Fischer et al., 2022). Our results are concordant with this observation as they show a similar pattern of improved protein performance using a diverse set of radii.

While the LGBM shows the overall best performance, there are tradeoffs to consider when choosing a machine learning model for single-cell protein abundance imputation in MTI datasets. Traditional ML models such as LGBM and EN can only impute one protein per model, which requires training and storing a model for each protein to be imputed. Using a single model for each protein is time and cost inefficient. In contrast, an AE can impute multiple proteins at once and even all proteins included in their training data, requiring only a single training session and model. While AEs perform marginally worse than LGBM and EN for protein imputation, their capability for multi-protein imputation offers an advantage in reduced training time and cost. Multi-protein imputation, as opposed to sequential imputation, also models inter-protein relationships, and potentially yields more biologically pertinent relationships to explore.

We have demonstrated robust performance of our imputation methods and biological significance of imputed values. Using an independent MTI dataset from a cohort of 26 breast cancers that included all major subtypes of the disease, our imputation methods showed similar performance to that in our primary cohort. Imputation results across these datasets suggest that our machine learning methods are versatile and can potentially be used in other MTI datasets. To demonstrate the biological significance of our imputed data, we used imputed protein abundance levels to accurately predict whether individual cells are taken from pre-treatment or post-treatment biopsies. This application shows that our approach produces biologically relevant imputed data that can be used in biomedical research.

Limitations of this work include a focus on protein abundance rather than RNA expression, the small number of proteins used for imputation, the cohort composition of metastatic breast cancers, and the small sample size. These analyses demonstrate that it is possible to impute protein abundance in MTI, but imputation of RNA expression has not been explored. This analysis used the sixteen proteins that were shared amongst all biopsies, and it is uncertain if other proteins can be imputed as accurately as these sixteen. This analysis also focused on breast cancer biopsies and diseased tissue, and imputation results may be different in healthy tissue or in other diseases. A study like ours would benefit from using a larger and more diverse cohort and from different MTI assays with more proteins. Different and larger datasets would help establish the robustness and generalizability of our imputation methods.

In summary, this study demonstrates that machine learning can effectively impute biologically meaningful single-cell protein abundance levels using MTI datasets. Our results provide a foundation for future applications of machine-learning imputed data in single-cell MTI datasets. One potential application is imputation of additional single-cell and cellular neighborhood features, which can in turn aid in understanding tissue ecosystems. Another future application is the use of imputed datasets to predict biomedical outcomes such as tissue response to perturbations or, in the case of disease, response to therapy.

## Data availability

The datasets used for this study are available in DataVerse. https://dataverse.harvard.edu/dataset.xhtml?persistentId=doi:10.7910/DVN/RBIJSQ&version=1.0 and Table 4 lists the HTAN Biopsy and Biospecimen IDs used in this study. More information on these biopsies and biospecimens is available at https://humantumoratlas.org/explore

**Table 4:**
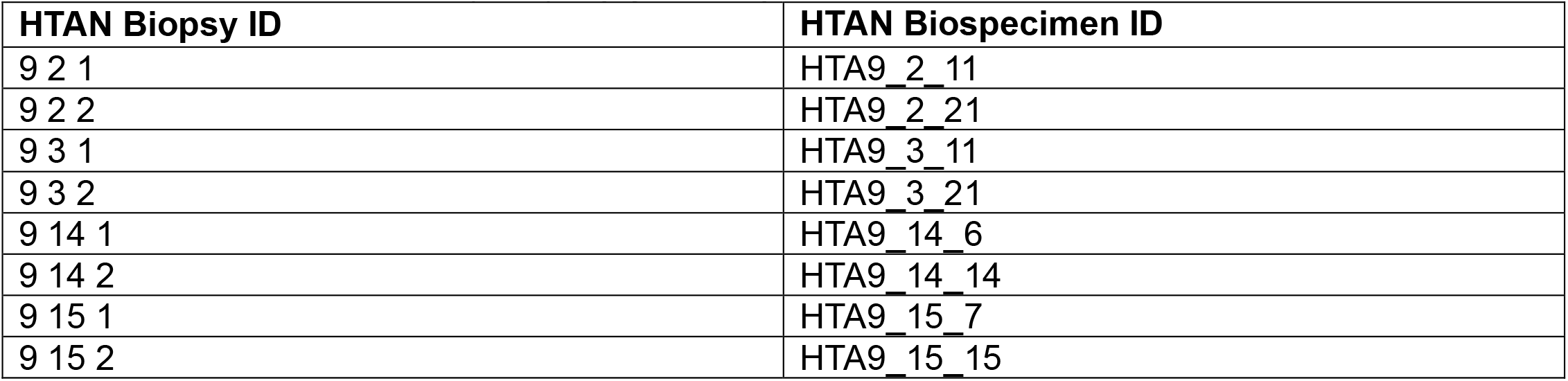
Human Tumor Atlas Network (HTAN) biopsy and biospecimen IDs.

All TMA data is available through the HTAN Data Portal as part of the HTAN TNP-TMA Project (https://data.humantumoratlas.org/)

## Code availability

The source code of this work is freely available in the GitHub repository. https://github.com/goeckslab/MTIProteinImputation

## Methods

### Experimental Setup

The BOND RX Automated IHC/ISH Stainer was used to bake FFPE slides at 60°C for 30 minutes, to dewax the sections using the Bond Dewax solution at 72°C, and for antigen retrieval using Epitope Retrieval 1 (Leica™) solution at 100°C for 20 minutes. Slides underwent multiple cycles of antibody incubation, imaging, and fluorophore inactivation. All antibodies were incubated overnight at 4°C in the dark. Slides were stained with Hoechst 33342 for 10 minutes at room temperature in the dark following antibody incubation in every cycle. Coverslips were wet-mounted using 200 μL of 10% Glycerol in PBS prior to imaging. Images were acquired using a 20x objective (0.75 NA) on a CyteFinder slide scanning fluorescence microscope (RareCyte Inc. Seattle WA). Fluorophores were inactivated using a 4.5% H2O2, 24 mM NaOH/PBS solution and an LED light source for 1 hour.

The detailed protocol is available in protocols.io (dx.doi.org/10.17504/protocols.io.bjiukkew).

### Data Preparation

The original source files include X and Y spatial coordinates and bio-morphological information (orientation, area, extent, etc.) for each cell. These features are removed for the initial imputation experiments, which solely rely on protein information.

To prepare the available data for the machine learning models and deep learning networks, we used Min-Max Scaling to scale features to be in the [0,1] range.

#### Statistical Validity

For robust statistical validity, we conducted more than 30 experiments (n > 30) for each protein imputation and each model.

### Elastic Net

To setup an experiment using the Elastic Net, the scikit-learn library was used and within this library the ElasticNetCV, which automatically performs cross validation of error values. To support our results with statistical significance as well as a high enough numbers of trials, we performed multiple experiments n > 30. Each run was performed using a different random seed.

### Light GBM

To set up a training and evaluation pipeline for our Light GBM (Ke et al., 2017) model, we used the Ludwig (Molino et al., 2019) platform, which enables “End-to-end machine learning pipelines” in a low code environment. To set up a Ludwig network only a config file specifying the features, in this work the proteins, and the target to be imputed is required. We automated this process by using a combination of shell and make scripts. Each run was assigned a different random seed to ensure reproducibility.

### Auto Encoder

The Auto Encoder imputation was set up to make use of an iterative approach. As a first step, the preprocessed source data was loaded. To impute a specific protein, the protein values were replaced with a mean value calculated by using all available values for the protein in question. After the dataset was prepared and the protein in question replaced, the data was used as input for the auto encoder. A full encode and decode process was performed (**Fig. 2**), and the output was stored as an intermediate result. From this intermediate result, the imputed protein values were taken and used as a replacement of the mean protein values created in the preparation step. After this, another round of imputation was performed. This process was preformed 10 times, which resulted in a 10-step iterative imputation process. Reported MAE and RMSE values are calculated by using the last 5 iterative decoding’s, calculating the mean of the decoding’s for the protein and then calculate the MAE and RMSE.

For each model and network, a minimum of 30 experiments were performed to establish statistical significance. To create reproducible but different results, each experiment used a unique random seed.

## Acknowledgements

This research was supported by the National Cancer Institute (NCI) of the National Institutes of Health grants U24CA231877 and U2CCA233280 and by funding from the Prospect Creek Foundation to the OHSU SMMART (Serial Measurement of Molecular and Architectural Responses to Therapy) Program.

## Contributions

R.K and J.G both conceived the study. C.W. and A.C. processed the primary image data to produce the single-cell datasets analyzed in this work. R.K. developed the methods, wrote the code, and performed the analysis. C.W., A.C., and J.G. provided feedback and suggestions on methods, code, and analyses. K.K. designed illustrations used in the figures and provided feedback on the overall figure design. All authors read and approved the final paper.

## Ethic Declarations

This research strictly adheres to ethical standards in data collection, processing, and usage. All datasets used for training and testing our machine learning models are publicly available.

