## Supplements for "Imputing Single-Cell Protein Abundance in Multiplex Tissue Imaging"

**Table S 1: Mean and (Standard Deviation) of all observed radii as well as the baseline for LGBM, AE Single and Multi-Imputation using an Across-Patient Setup.**

| Network | 0 µm/ Baseline | 15 µm | 30 µm | 60 µm | 90 µm | 120 µm |
|---|---|---|---|---|---|---|
| LGBM | 0.099 (0.060) | 0.101 (0.059) | 0.101 (0.059) | 0.097 (0.055) | 0.104 (0.062) | 0.106 (0.068) |
| AE Single Protein | 0.128 (0.089) | 0.100 (0.064) | 0.105 (0.059) | 0.113 (0.062) | 0.120 (0.076) | 0.119 (0.075) |
| AE Multi Protein | 0.120 (0.087) | 0.115 (0.068) | 0.112 (0.062) | 0.117 (0.068) | 0.120 (0.081) | 0.121 (0.081) |

**Table S 2: Variance for each observed protein.**

| Protein | Variance |
|---|---|
| CK19 | 1.28 |
| Vimentin | 0.42 |
| ER | 0.32 |
| pERK | 0.24 |
| aSMA | 0.22 |
| pRB | 0.18 |
| Ecad | 0.14 |
| AR | 0.13 |
| CK14 | 0.11 |
| CD45 | 0.09 |
| HER2 | 0.09 |
| Ki67 | 0.08 |
| EGFR | 0.06 |
| P21 | 0.05 |
| PR | 0.05 |
| CK17 | 0.05 |

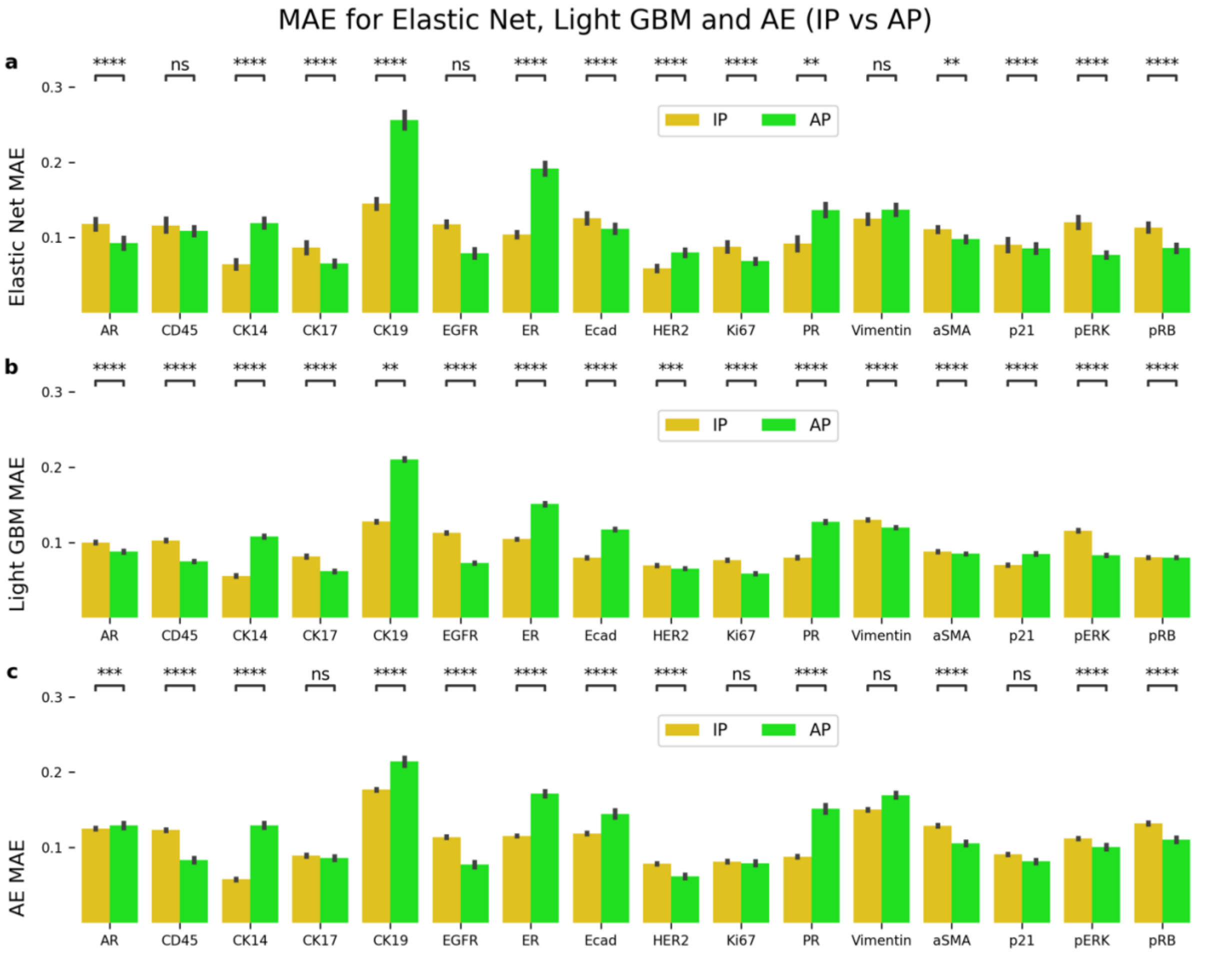


**Figure S 1: Performance comparison between In Patient (IP) and Across Patient (AP).** In Patient (IP, golden) is compared to Across Patient (AP, green) across models: **a**) Elastic Net (top), **b**) Light GBM (middle) and **c**) Autoencoder (AE, bottom). For means of comparison the Mean Absolute Error (MAE) was calculated. Statistical analysis using Mann-Whittney-Wilcoxon and multi-hypothesis testing using Benjamini-Hochberg correction. p-values: ns: not significant, p <= 1.00e+00 *: 1.00e-02 < p <= 5.00e-02 **: 1.00e-03 < p <= 1.00e-02 ***: 1.00e-04 < p <= 1.00e-03 ****: p <= 1.00e-0

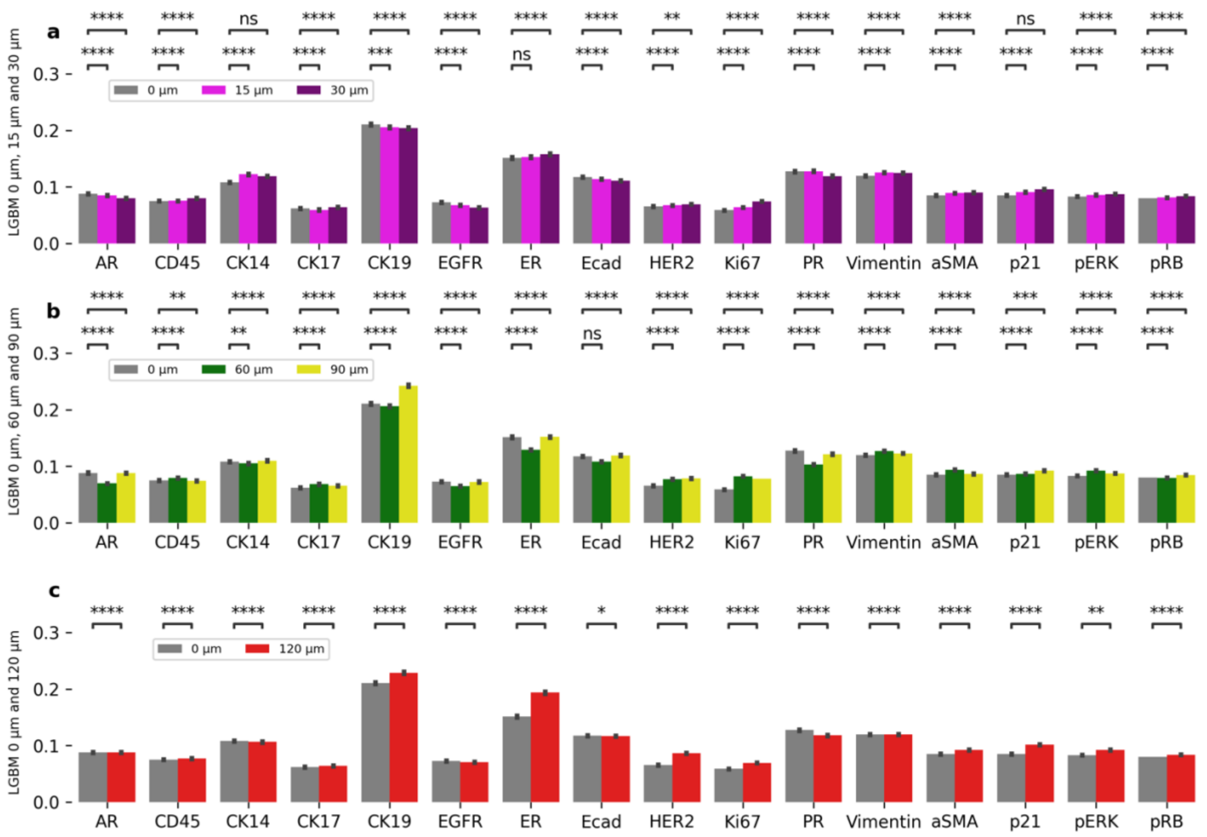


**Figure S 2: Performance comparison between baseline (0 µm) and increasing spatial distances of 15, 30, 60, 90 and 120 µm.** Light GBM results comparing increasing spatial distances to the baseline (0 µm in grey) across proteins. **a)** Baseline compared to 15 µm (magenta) and 30 µm (plum). **b)** Baseline compared to 60 µm (green) and 90 µm (yellow) and **c)** Baseline compared to 120 µm (red). For means of comparison the Mean Absolute Error (MAE) was calculated. Statistical analysis using Mann-Whittney-Wilcoxon and multi-hypothesis testing using Benjamini-Hochberg correction. p-values: ns: not significant, p <= 1.00e+00 *: 1.00e-02 < p <= 5.00e-02 **: 1.00e-03 < p <= 1.00e-02 ***: 1.00e-04 < p <= 1.00e-03 ****: p <= 1.00e-0-

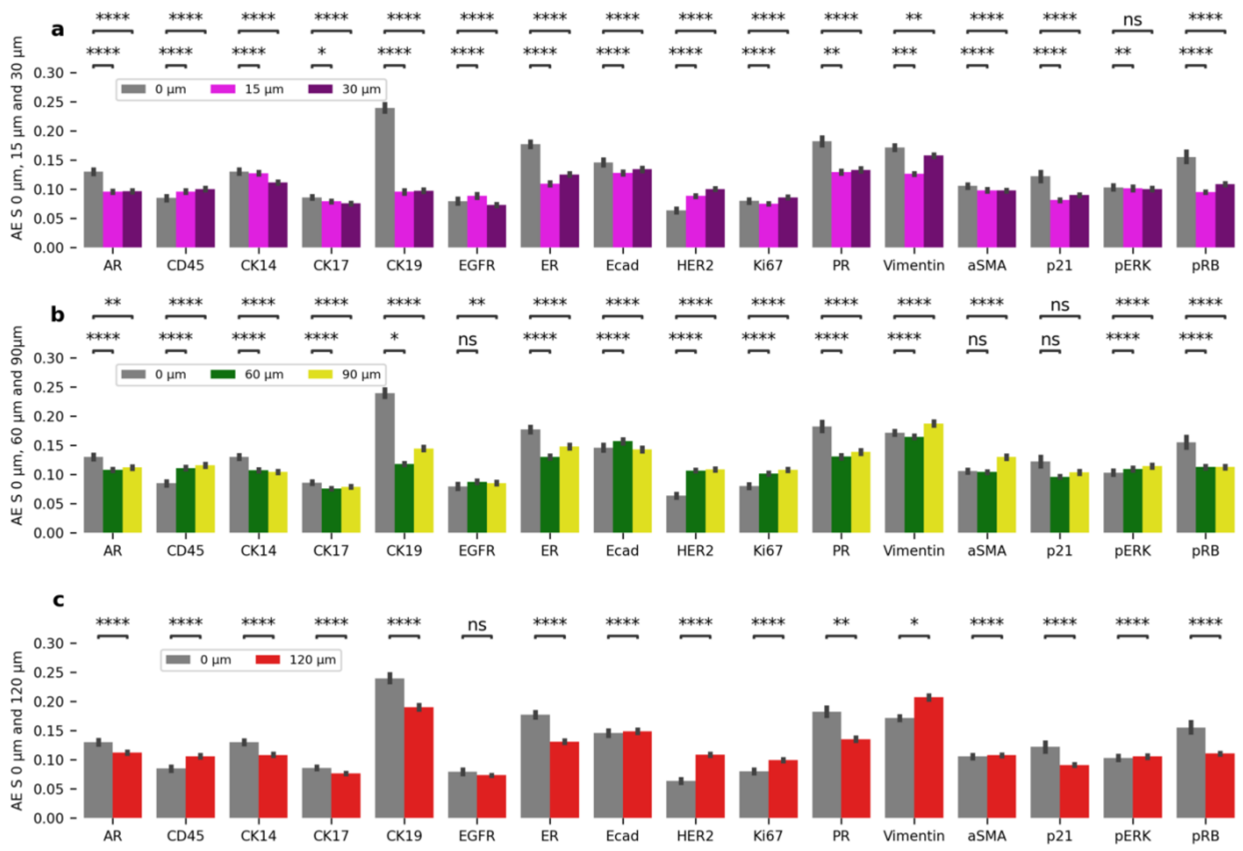


**Figure S 3: Performance comparison between baseline (0 µm) and increasing spatial distances of 15, 30, 60, 90 and 120 µm.** Autoencoder (AE) results comparing increasing spatial distances to the baseline (0 µm in grey) across proteins. **a)** Baseline compared to 15 µm (magenta) and 30 µm (plum). **b)** Baseline compared to 60 µm (green) and 90 µm (yellow) and **c)** Baseline compared to 120 µm (red). For means of comparison the Mean Absolute Error (MAE) was calculated. Statistical analysis using Mann-Whittney-Wilcoxon and multi-hypothesis testing using Benjamini-Hochberg correction. p-values: ns: not significant, p <= 1.00e+00 *: 1.00e-02 < p <= 5.00e-02 **: 1.00e-03 < p <= 1.00e-02 ***: 1.00e-04 < p <= 1.00e-03 ****: p <= 1.00e-0

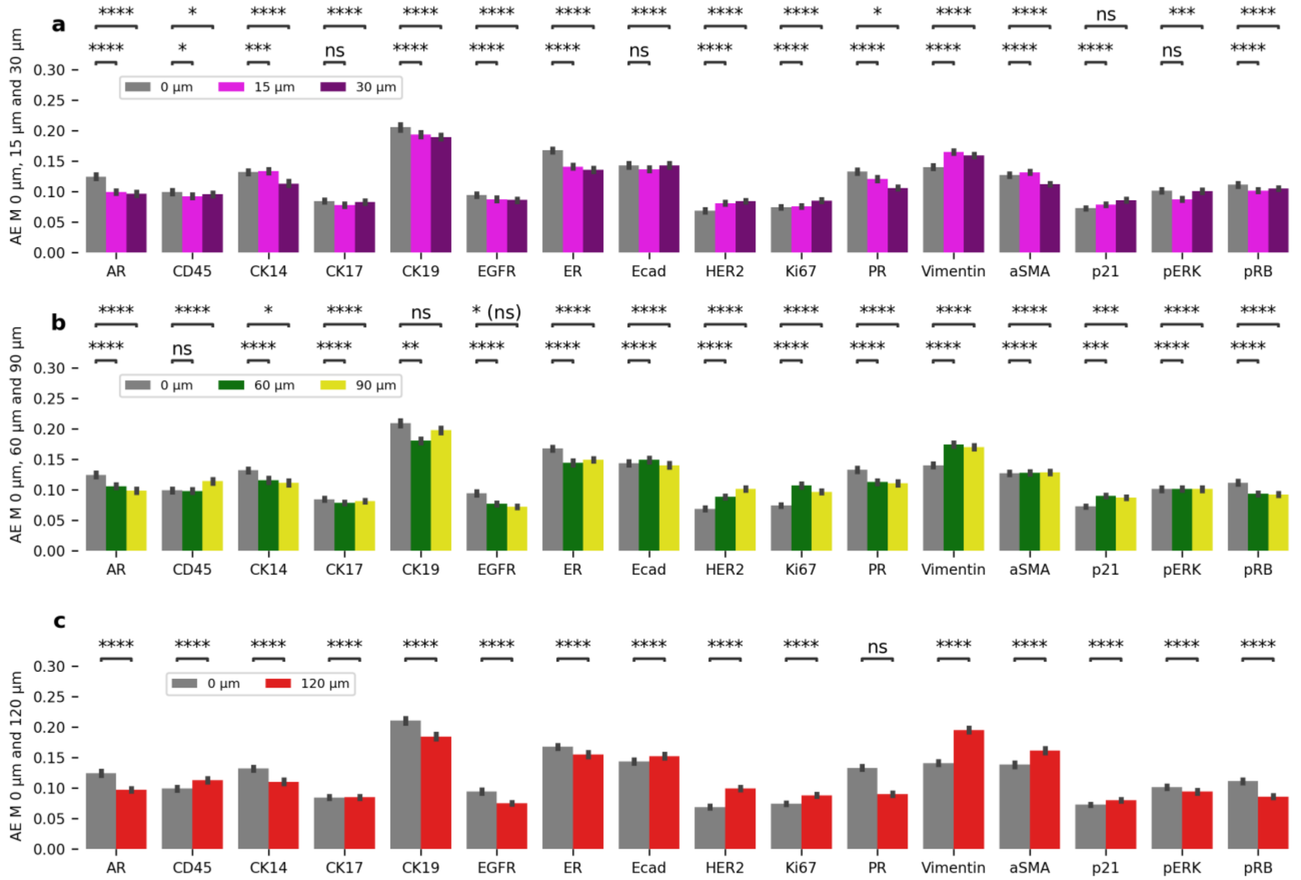


**Figure S 4: Performance comparison between baseline (0 µm) and increasing distances of 15, 30, 60, 90 and 120 µm.** Autoencoder (AE) results using multi protein imputation comparing increasing spatial distances to the baseline (0 µm in grey) across proteins. **a)** Baseline compared to 15 µm (magenta) and 30 µm (plum). **b)** Baseline compared to 60 µm (green) and 90 µm (yellow) and **c)** Baseline compared to 120 µm (red). For means of comparison the Mean Absolute Error (MAE) was calculated. Statistical analysis using Mann-Whittney-Wilcoxon and multi-hypothesis testing using Benjamini-Hochberg correction. p-values: ns: not significant, p <= 1.00e+00 *: 1.00e-02 < p <= 5.00e-02 **: 1.00e-03 < p <= 1.00e-02 ***: 1.00e-04 < p <= 1.00e-03 ****: p <= 1.00e-0
